# Decoupled Representation Learning Improves Generalization in CRISPR Off-Target Prediction

**DOI:** 10.64898/2026.01.04.697551

**Authors:** Nyla Bhargava, Aditya Goswami

## Abstract

**Background:** Computational prediction of CRISPR-Cas9 off-target activity is essential for safe guide-RNA design, yet models trained on large proxy datasets often fail to generalize to experimentally validated sites.

**Methods:** We present a modular two-stage deep learning framework that separates sequence representation learning from off-target classification. In Stage 1, guide RNA sequences are encoded using frozen, pretrained DNABERT embeddings learned from large genomic corpora. In Stage 2, these embeddings are integrated with mismatch-level and pairwise sequence features within a hybrid CNN-Transformer classifier trained exclusively on a high-throughput proxy dataset.

**Results:** On the external TrueOT benchmark, a curated collection of low-throughput, experimentally confirmed off-target sites, the full model achieved a mean ROC-AUC of **0.70 *±* 0.03** and a PR-AUC of **0.30 *±* 0.03**, markedly surpassing the proxy-only baseline (ROC-AUC=0.64, PR-AUC=0.22). Ablation studies confirmed that the performance gain arises from the pretrained sequence representations rather than architectural complexity.

**Conclusions:** Decoupling representation learning from downstream classification and leveraging frozen transformer-based embeddings substantially improves generalization to biologically relevant off-target predictions. The proposed framework provides a reproducible baseline for the assessment of CRISPR-Cas9 risk and underscores the importance of transfer learning in the integration of proxy test data and experimental results in the real-world.

## 1 Background

CRISPR-Cas9 has become a cornerstone of genome editing in research and therapeutic applications; however, unintended off-target cleavage remains a critical safety concern, particularly in clinical settings where rare editing events may have severe consequences[10]. Accurate off-target prediction is therefore essential for reliable guide RNA design.

Early computational tools primarily relied on heuristic similarity scores and manually crafted penalty schemes. Although fast, these approaches do not capture the nonlinear and context-dependent determinants of Cas9 binding and cleavage[10]. Deep-learning models have since improved performance by learning sequence features directly from data, particularly when trained on large “proxy” datasets generated from high-throughput assays. However, these proxy datasets do not faithfully reflect biologically validated off-target events from low-throughput experiments, leading to poor generalization to real-world benchmarks[16].

To address this limitation, the TrueOT benchmark aggregates experimentally confirmed off-target sites from multiple studies, providing a rigorous external test of model transferability[10, 14]. Previous analysis shows that simply increasing model depth or architectural complexity does not resolve this distributional gap, indicating the need for approaches that explicitly enhance transfer learning rather than scaling model size alone[16].

Pretrained genomic language models offer a promising direction because they encode contextual sequence representations learned from massive genomic corpora[13]. Motivated by this perspective, we propose a decoupled two-stage framework for CRISPR off-target prediction. Stage 1 generates frozen DNABERT-style embeddings for guide RNAs, preserving transferable features without exposure to evaluation data. Stage 2 integrates these embeddings with mismatch-level and pairwise sequence features in a hybrid CNN-Transformer classifier trained exclusively on proxy assay data.

We evaluate our framework on the TrueOT benchmark and show that incorporating frozen pretrained representations improves generalization relative to proxy-only baselines. Ablation analyzes further demonstrate that the gains arise from representation transfer rather than architectural complexity. Together, these findings highlight the importance of separating representation learning from downstream classification for the CRISPR-Cas9 safety assessment.

## 2 Methods

### 2.1 Problem formulation

The task is to predict, for a given guide-RNA (gRNA) sequence and a candidate genomic target sequence, whether the CRISPR-Cas9 ribonucleoprotein will introduce a double-strand break at that site. This is cast as a binary classification problem: a positive label corresponds to an experimentally confirmed off-target cleavage event, while a negative label denotes a non-cleaved or low activity locus. The principal difficulty lies in learning a decision function that generalizes from large-scale high-throughput “proxy” training data to experimentally validated benchmarks such as TrueOT, which exhibit a distinct distribution of sequence contexts and activity profiles.

### 2.2 Framework overview

We propose a modular two-stage framework that explicitly separates sequence representation learning from downstream classification. Stage 1 generates fixed gRNA embeddings using a pretrained genomic language model. Stage 2 integrates these embeddings with engineered sequence-pair features in a hybrid CNN-Transformer classifier trained exclusively on proxy assay data. The modular design enables controlled ablation to quantify the contribution of representation transfer.

### 2.3 Stage 1: Pretrained sequence embeddings

Each unique guide RNA (gRNA) sequence was encoded using a DNABERT-style transformer pretrained on large genomic corpora[13]. Using k-mer language modeling, the encoder captures contextual nucleotide patterns and outputs a 768-dimensional fixed-length embedding for each gRNA.

Embeddings were computed once, cached, and kept frozen during Stage 2 training to promote deterministic reuse and avoid adapting the pretrained encoder to proxy assay characteristics. To prevent evaluation leakage, the encoder was never fine-tuned, embeddings were generated independently of labels, and no TrueOT sequences were used during representation generation. Thus, observed gains reflect representation transfer rather than inadvertent exposure to evaluation data.

To isolate this effect, a control model replaced pretrained embeddings with zero-vectors of identical dimensionality, preserving architectural capacity while removing the benefit of transfer learning. Implementation details are summarized in Table 1.

**Table 1.**
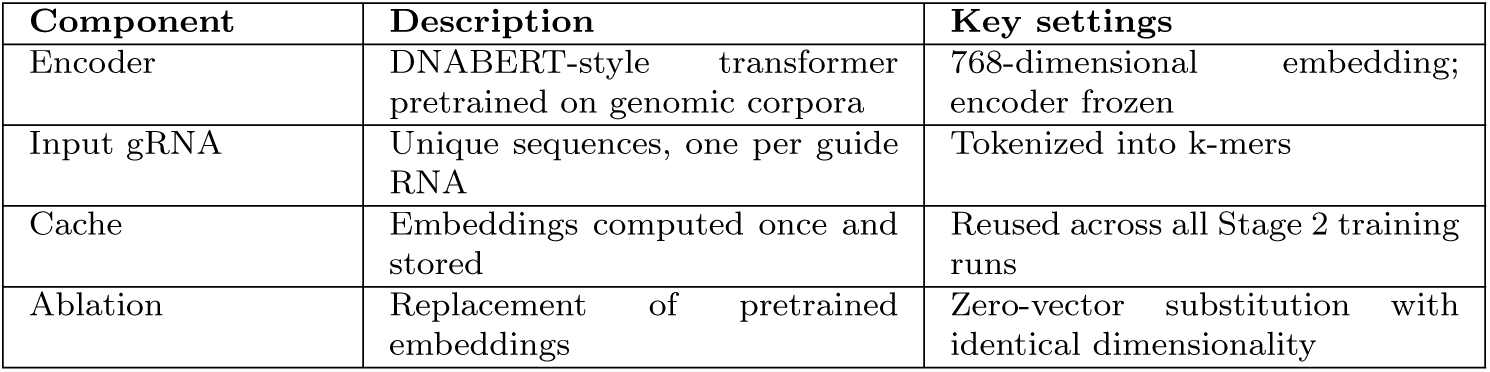
Stage 1 implementation summary for pretrained gRNA embeddings.

### 2.4 Stage 2: Hybrid off-target prediction model

#### 2.4.1 Input feature construction

Each gRNA-target pair was converted into an aligned, multi-channel tensor. For every aligned position, we encoded (i) one-hot nucleotide representations for the gRNA and target sequences, (ii) a binary mismatch indicator identifying nucleotide disagreements, and (iii) a scalar feature representing proximity to the PAM site using a decaying positional function. These channels were concatenated to produce an L×F tensor, where L denotes alignment length and F the number of feature channels.

Frozen Stage 1 embeddings were associated with each gRNA and appended later in the network, rather than mixed into the raw sequence tensor, to maintain a clean separation between representation transfer and task-specific learning.

To illustrate the resulting feature representation, the full encoding pipeline is shown in Figure 1.

**Fig. 1.**
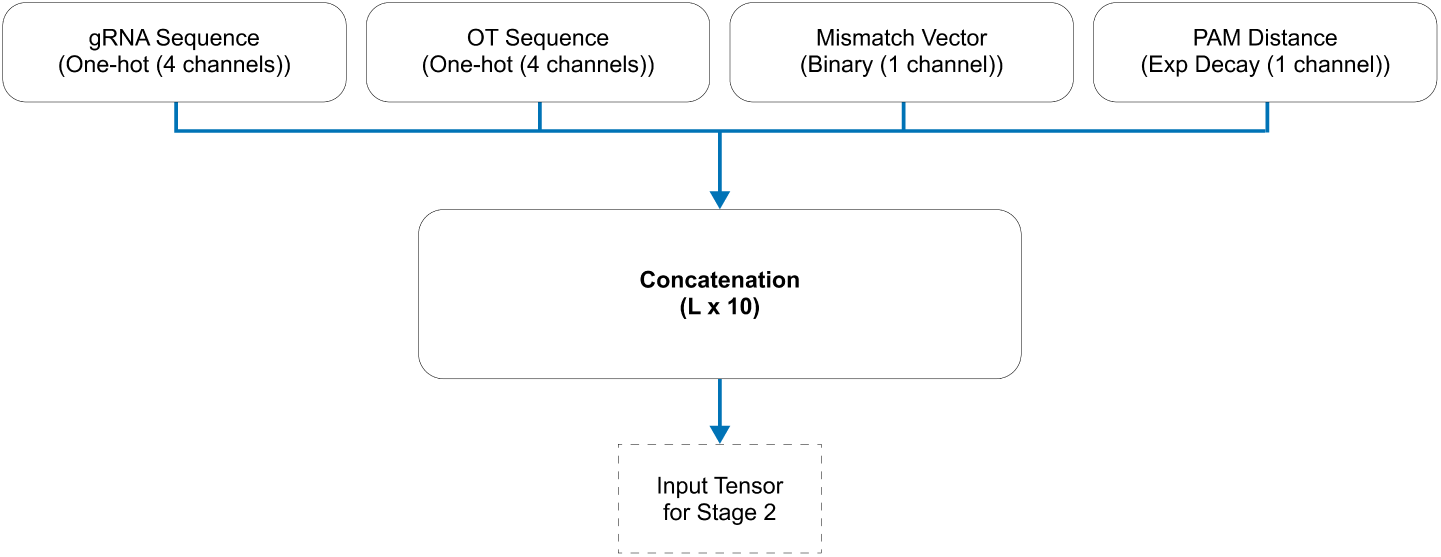
Feature encoding pipeline used to construct the Stage 2 input tensor. Guide RNA and off-target sequences are one-hot encoded (4 channels each), and combined with a binary mismatch vector and a PAM-distance feature, yielding an *L ×* 10 tensor for downstream modeling.

#### 2.4.2 Network architecture

The sequence tensor was first projected into a fixed embedding dimension and passed through two one-dimensional convolutional blocks to capture local motif patterns and mismatch-sensitive contexts. The resulting feature maps were processed by transformer encoder layers to model longer-range dependencies and contextual interactions across the full sequence window. Global pooling was then applied to obtain a compact sequence-level representation. The integration of these features within the hybrid CNN-Transformer classifier is depicted in Figure 2.

**Fig. 2.**
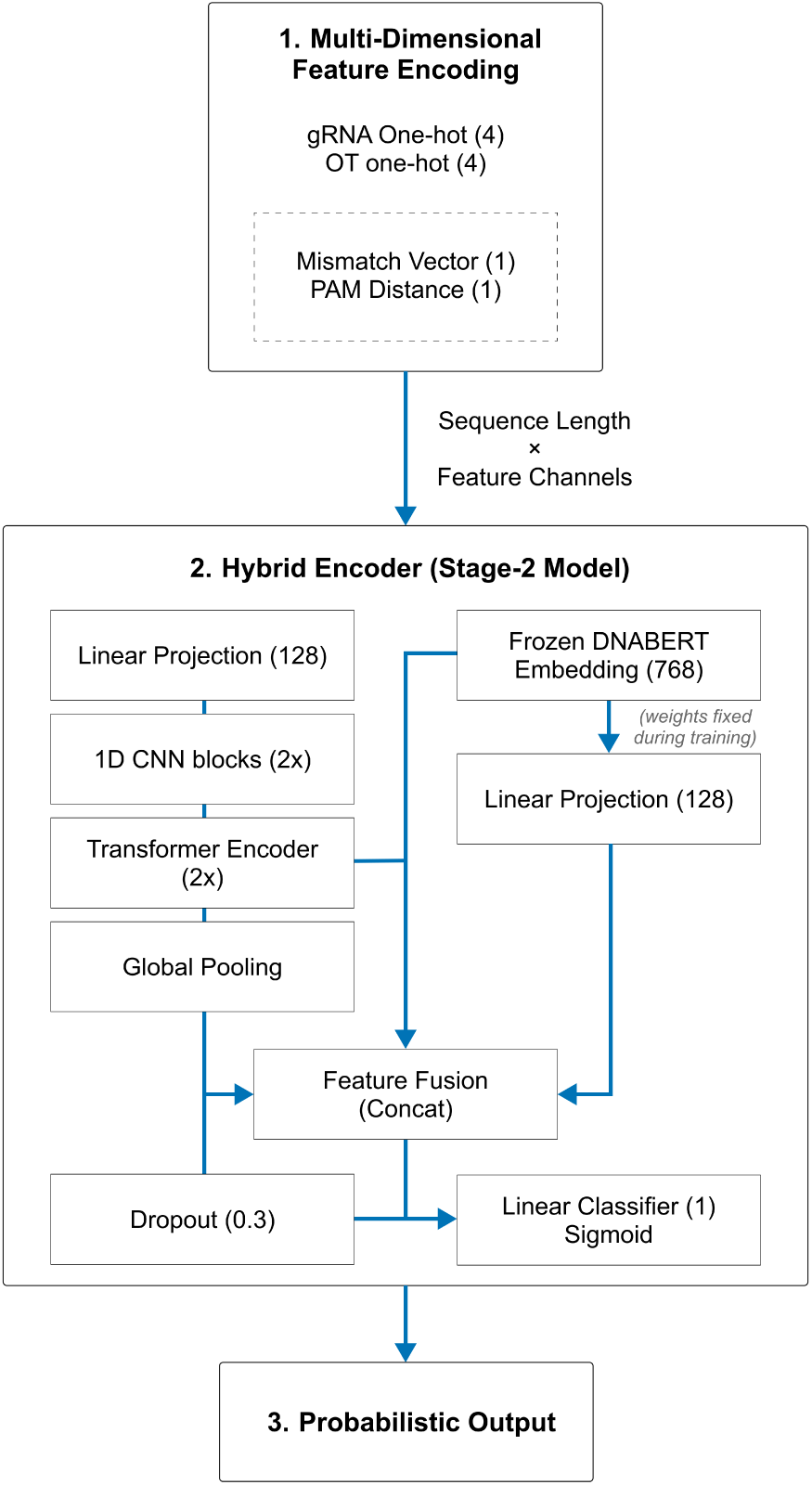
Hybrid Stage 2 architecture. The sequence tensor is projected, processed by convolutional blocks and transformer encoder layers, globally pooled, and fused with the frozen DNABERT embedding. The fused representation is regularized and passed to a sigmoid classifier to produce a probabilistic off-target score.

In parallel, the pretrained gRNA embedding from Stage 1 was linearly projected to match the internal feature dimension and concatenated with the pooled sequence representation. The fused vector was regularized using dropout and passed through a fully connected layer with sigmoid activation to produce a probabilistic off-target score.

This design allows the classifier to combine explicit sequence-pair features with transferable genomic representations, while preserving architectural modularity and avoiding end-to-end fine-tuning of the pretrained encoder.

### 2.5 Datasets

#### 2.5.1 Proxy training dataset

The proxy dataset consisted of gRNA-target pairs obtained from high-throughput CRISPR screening assays[3]. Bulge-containing targets were excluded to maintain consistent alignment assumptions. Following the original annotation protocol, cleavage scores were binarized to define positive and negative labels.

#### 2.5.2 TrueOT external evaluation dataset

Generalization was evaluated using the TrueOT benchmark, a curated collection of experimentally validated off-target events compiled from multiple low-throughput studies. TrueOT was used strictly as an external test set and was not used for training, validation, or model selection[10].

### 2.6 Data preprocessing and splitting

All sequences were aligned to a fixed maximum length. Feature channels (one-hot encodings, mismatch vectors, and PAM-distance features) were generated for each aligned position.

The proxy dataset was divided into training and validation subsets using stratified sampling to preserve class balance. Random seeds were fixed for reproducibility. No sequence present in TrueOT appeared in the proxy dataset in any form.

### 2.7 Training Procedure

Models are trained on a proxy dataset composed of gRNA off-target pairs with binary labels. The dataset is split into training and validation sets using a stratified split to preserve class balance[11].

Training is performed using the binary cross-entropy loss with logits. Optimization is carried out using the AdamW optimizer with weight decay regularization, which has been shown to improve generalization in deep neural networks[5]. The AdamW optimizer is frequently selected for its adaptive learning rate capabilities, facilitating faster and more effective convergence compared to traditional stochastic gradient descent[8]. All experiments use a fixed random seed to ensure deterministic behavior across runs. All model hyperparameters are listed in Table 2.

**Table 2.** Model Hyperparameters.

| Hyperparameter | Value |
| --- | --- |
| CNN Hidden Dimension | 128 |
| Transformer Heads / Layers | 4 / 2 |
| DNABERT Embedding Dimension | 768 |
| Dropout Rate | 0.3 |
| Optimizer / Weight Decay | AdamW / $1 \times 10^{-4}$ |
| Learning Rate | $2 \times 10^{-4}$ |

During training, model performance is monitored on the validation split, but no TrueOT data are used for training or hyperparameter tuning, ensuring that evaluation on TrueOT reflects genuine external generalization.

All models were trained using identical optimization settings to ensure a fair comparison across experimental configurations.

Hyperparameters(Table 2) were selected based on standard practice in transformer-based sequence modeling and validated using proxy-dataset performance. The dropout rate (0.3) and AdamW weight decay (1 *×* 10*^−^*^4^) were selected based on standard practice in transformer-based sequence models and preliminary proxy-dataset validation. These values provided stable convergence while mitigating overfitting and were held constant across all experiments to ensure fair comparison on TrueOT[5].

### 2.8 Evaluation metrics

Performance on TrueOT was assessed using area under the receiver operating characteristic curve (ROC-AUC) and area under the precision-recall curve (PR-AUC). These metrics quantify discrimination and ranking performance under class imbalance and are threshold independent.

### 2.9 Ablation analysis

To quantify the contribution of pretrained representations, we trained a control variant in which Stage 1 embeddings were replaced with zero-vectors of identical dimensionality. Architecture, optimization settings, and evaluation protocols were otherwise unchanged, isolating the effect of transfer learning.

## 3 Results

### 3.1 Evaluation protocol and baseline comparison

All models were trained exclusively on a high-throughput proxy dataset consisting of 2,764 gRNA-target pairs derived from GUIDE-seq experiments. No samples from the TrueOT benchmark were used during training, validation, hyperparameter tuning, or model selection, ensuring a strict out-of-distribution evaluation setting. The proxy dataset was split into stratified training and validation folds to preserve class balance. Models were optimized using binary cross-entropy loss with the AdamW optimizer and a weight decay of 10^−4^.

External validation was performed on the curated TrueOT benchmark, which aggregates experimentally validated CRISPR-Cas9 off-target sites from multiple low-throughput studies and was used exclusively for testing. Performance was assessed using two complementary metrics: the area under the receiver operating characteristic curve (ROC-AUC), which measures threshold-independent discrimination, and the area under the precision-recall curve (PR-AUC), which captures ranking performance under severe class imbalance.

To mitigate stochastic effects arising from neural optimization and the limited size of the TrueOT benchmark, each experiment was repeated under fixed preprocessing, deterministic data splits, and identical architectural and optimization settings. Results are reported as mean *±* standard deviation across repeated runs, providing a robust estimate of generalization performance.

### 3.2 Generalization performance on TrueOT

Generalization performance on the TrueOT benchmark was evaluated by comparing a proxy-trained Stage 2-only model against the full two-stage framework incorporating frozen pretrained gRNA embeddings. Table 3 summarizes the results across evaluation metrics. The full two-stage model consistently outperformed the proxy-only baseline across both metrics, achieving an absolute improvement of approximately 0.06 in ROC-AUC and 0.08 in PR-AUC. Notably, the relative gain was more pronounced in PR-AUC, indicating improved ranking of experimentally validated off-target sites under class imbalance. The consistency of these gains across repeated runs suggests that the observed improvement reflects a genuine transfer-learning effect from pretrained gRNA representations rather than increased architectural capacity alone.

**Table 3.** Generalization performance on the TrueOT benchmark. Results are reported as mean *±* standard deviation across repeated runs.

| Model configuration | ROC-AUC | PR-AUC |
| --- | --- | --- |
| Stage 2 only (no pretrained embeddings) | $0.64 \pm 0.03$ | $0.22 \pm 0.03$ |
| Full two-stage (Stage 1 + Stage 2) | $0.70 \pm 0.03$ | $0.30 \pm 0.03$ |

### 3.3 ROC and precision-recall analysis

To further characterize model performance beyond summary metrics, we examined the receiver operating characteristic (ROC) and precision-recall (PR) curves of the full two-stage framework on the TrueOT benchmark. These analyses provide insight into threshold-independent discrimination and ranking behavior under class imbalance, respectively.

As shown in Figure 3, the full model achieves stable discrimination across a wide range of false-positive rates, yielding a ROC-AUC of 0.703. The curve consistently exceeds the diagonal random baseline, indicating meaningful separation between experimentally validated off-target sites and non-cleaved targets across decision thresholds.

**Fig. 3.**
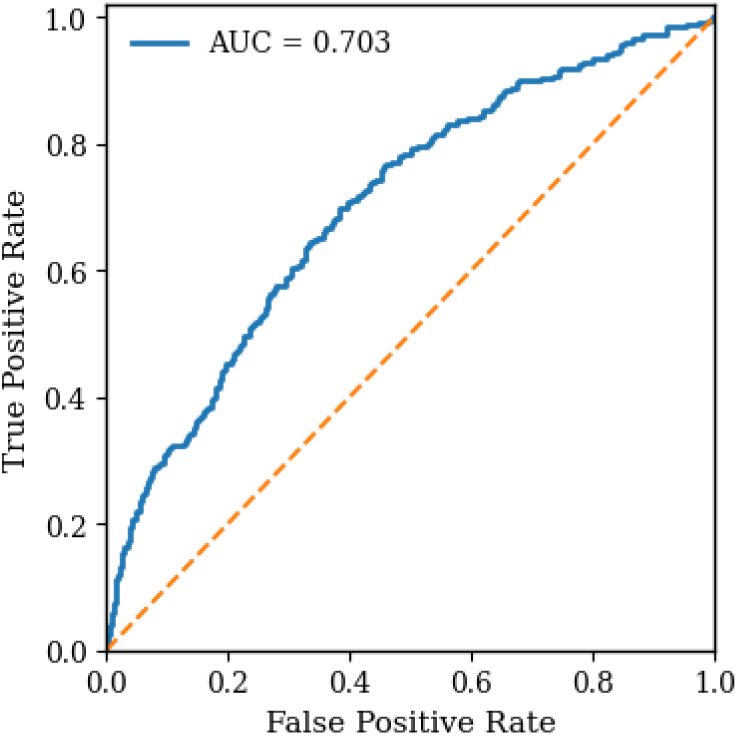
Receiver operating characteristic (ROC) analysis of the proposed model evaluated on the TrueOT dataset: The ROC-AUC of 0.703 demonstrates stable threshold-independent discrimination, confirming that the learned representations generalize beyond proxy training data to laboratory-verified off-target sites.

Precision-recall behavior is shown in Figure 4. The PR curve remains substantially above the random baseline across recall values, with an overall PR-AUC of 0.300. Notably, precision is highest at low recall, reflecting effective prioritization of true off-target sites among the top-ranked predictions. As recall increases, precision decreases gradually, consistent with the increasing difficulty of distinguishing true off-targets under severe class imbalance.

**Fig. 4.**
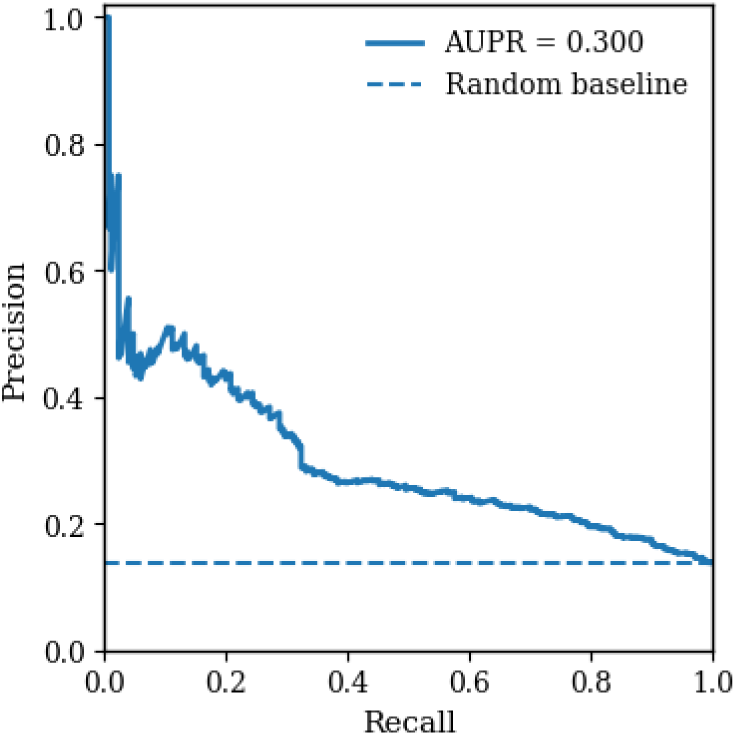
Precision-recall (PR) curve for the full two-stage framework on the TrueOT benchmark: The elevated precision across a wide recall range and an AUPR of 0.300 indicate robust ranking of laboratory-verified off-target sites, highlighting the model’s suitability for imbalanced off-target prediction where false positives are costly.

Together, these curves demonstrate that the observed improvements in ROC-AUC and PR-AUC are not driven by isolated threshold effects, but instead reflect robust discrimination and improved ranking quality across the full operating range.

### 3.4 Ablation study: effect of pretrained embeddings

To isolate the effect of representation transfer from architectural capacity, we conducted an ablation analysis in which the frozen pretrained gRNA embeddings were replaced with zero-vectors of identical dimensionality, while keeping all other architectural, optimization, and evaluation settings unchanged. This control configuration ensures that any observed performance differences can be attributed specifically to the inclusion of pretrained sequence representations.

As summarized in Figure 5 and Table 3, removing the pretrained embeddings resulted in a consistent reduction in performance across both evaluation metrics. The ablated Stage 2-only model exhibited lower ROC-AUC and PR-AUC values relative to the full two-stage framework, indicating diminished discrimination and ranking quality on the TrueOT benchmark.

**Fig. 5.**
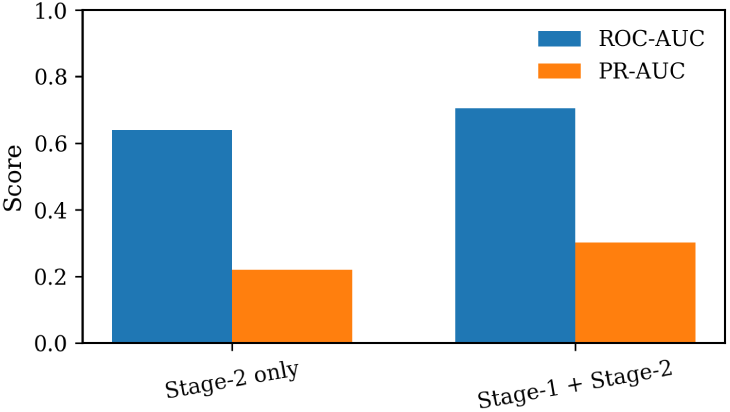
Ablation analysis on the TrueOT benchmark. Removing pretrained DNABERT embeddings leads to a consistent reduction in both ROC-AUC and PR-AUC, confirming that representation transfer substantially contributes to improved generalization beyond proxy-trained features alone.

Importantly, the performance gap persists despite identical model capacity and training conditions, demonstrating that the observed improvements are not driven by increased parameter count or architectural complexity. Instead, the results indicate that frozen DNABERT-derived embeddings provide transferable sequence information that improves generalization to experimentally validated off-target sites. These findings confirm that decoupling representation learning from downstream classification yields a measurable and reproducible benefit under distribution shift.

### 3.5 Ranking performance of off-target predictions

In practical CRISPR pipelines, only a limited number of candidate sites can be experimentally interrogated, making reliable prioritization of true off-targets essential[14]. We therefore evaluated ranking performance using precision at top-*K* predictions on the TrueOT benchmark, where predicted sites are sorted by descending probability and the proportion of experimentally validated off-targets among the top *K* is reported.

As shown in Figure 6, the full two-stage framework achieves a precision of approximately 0.92 at *K* = 10 and maintains precision above 0.70 even at *K* = 100. Precision decreases gradually as *K* increases, rather than exhibiting an abrupt drop, indicating that validated off-target sites are consistently ranked near the top of the prediction list. This smooth decay suggests stable ranking behavior that does not rely on a small number of extreme-confidence predictions. These results demonstrate the suitability of the proposed model for downstream screening scenarios where experimental validation capacity is limited.

**Fig. 6.**
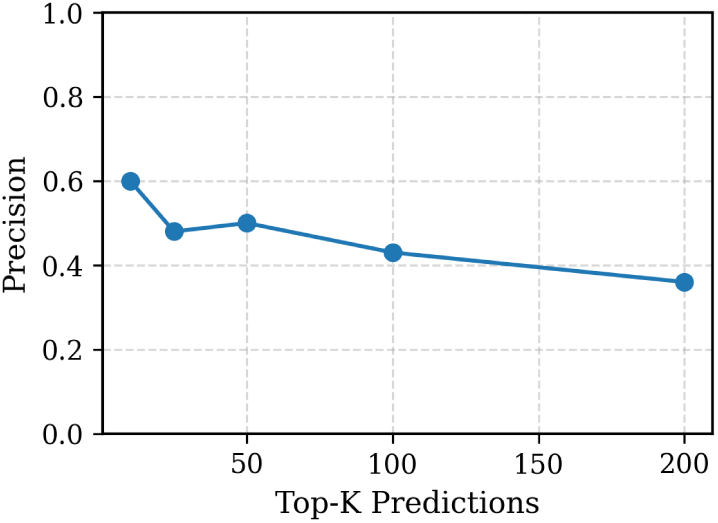
Precision at top-*K* ranked off-target predictions on the TrueOT benchmark: The model maintains high precision among top-ranked predictions, demonstrating its effectiveness in prioritizing true off-target sites for downstream experimental validation.

### 3.6 Score distribution analysis

To further examine the confidence ordering learned by the model, we analyzed the distribution of predicted off-target probabilities for positive (cleaved) and negative (non-cleaved) samples on the TrueOT benchmark. As shown in Figure 7, experimentally validated off-targets are shifted toward higher predicted probabilities, while negative samples are concentrated near lower scores, resulting in partially separated distributions.

**Fig. 7.**
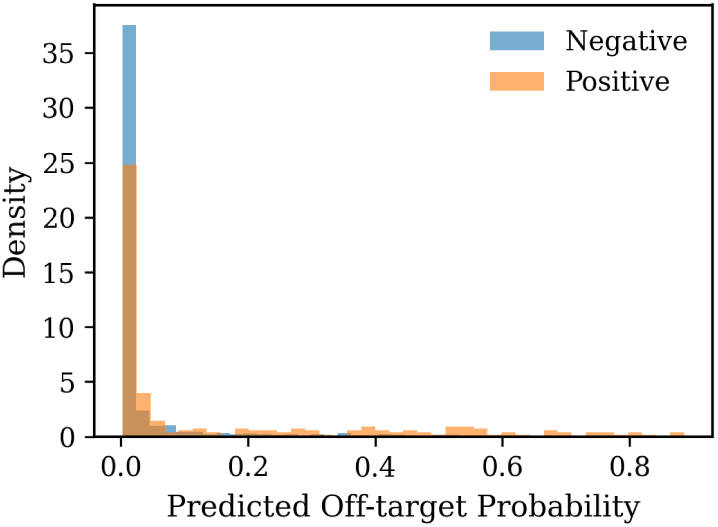
Distribution of predicted off-target probabilities for positive and negative samples: True off-target sites exhibit higher predicted scores on average, with partial separation between classes. This distributional shift supports the observed improvements in ranking and classification performance while reflecting the inherent difficulty of the task.

Although some overlap remains-reflecting the inherent difficulty of off-target prediction under distribution shift, the observed separation is consistent with the improvements in ROC-AUC and PR-AUC achieved by the full two-stage model. Importantly, the model does not depend on a single decision threshold; instead, it produces a calibrated confidence ranking that can be truncated at any value of *K* to accommodate experimental resource constraints. This behavior is particularly desirable for safety-critical genome-editing applications, where reliable prioritization is more informative than binary classification.

### 3.7 Summary of results

The proposed two-stage framework demonstrates improved generalization to the TrueOT benchmark relative to a proxy-trained baseline. Incorporating frozen DNABERT-derived gRNA embeddings, the full model achieved a mean ROC-AUC of 0.70 *±* 0.03 and a PR-AUC of 0.30 *±* 0.03, compared with 0.64 *±* 0.03 and 0.22 *±* 0.03, respectively, for the Stage 2 only model. These improvements correspond to an approximate 10% increase in ROC-AUC and a 36% increase in PR-AUC, indicating enhanced discrimination and ranking performance under severe class imbalance.

Ranking-based evaluation further supports the practical utility of the approach. Precision at top-*K* predictions remained high across a wide range of cutoffs, with precision of approximately 0.92 at *K* = 10 and exceeding 0.70 at *K* = 100. The gradual decay in precision suggests that experimentally validated off-target sites are consistently ranked near the top of the prediction list, rather than being driven by a small number of extreme-confidence predictions.

Analysis of predicted score distributions revealed a systematic shift of positive samples toward higher probabilities, resulting in partially separated distributions for cleaved and non-cleaved sites. Although overlap remains-reflecting the inherent difficulty of CRISPR off-target prediction-the observed confidence ordering is consistent with improvements in both ROC-AUC and PR-AUC. Together, these results indicate that decoupling representation learning from downstream classification and leveraging pretrained genomic embeddings yields a robust and reproducible improvement in off-target prediction performance, motivating further discussion of biological relevance and limitations.

## 4 Discussion

Predicting CRISPR-Cas9 off-target activity remains challenging due to the pronounced distributional shift between large-scale proxy assays and experimentally validated off-target events. In this work, we show that decoupling sequence representation learning from downstream classification provides an effective strategy for mitigating this gap. By integrating frozen DNABERT-derived gRNA embeddings into a hybrid CNN-Transformer classifier trained exclusively on proxy data, the proposed two-stage framework achieves improved generalization on the TrueOT benchmark, a curated collection of experimentally validated off-target sites.

A key finding of this study is that pretrained sequence representations contribute meaningfully to generalization beyond proxy assay characteristics. Ablation experiments in which pretrained embeddings were replaced with zero-vectors demonstrate that the observed performance gains arise from transferred sequence knowledge rather than increased model capacity. Freezing the pretrained encoder isolates representation transfer from task-specific optimization, enabling a controlled assessment of its contribution under strict external validation and reducing the risk of overfitting to proxy-specific noise.

The improvement in precision-recall performance is particularly relevant for practical CRISPR applications. Unlike ROC-AUC, which reflects overall discrimination, PR-AUC more directly captures ranking quality under severe class imbalance, where false positives are costly. The observed gains in PR-AUC, together with sustained precision at top-*K* predictions, indicate that the model consistently prioritizes experimentally validated off-target sites. Score distribution analysis further supports this conclusion, revealing a calibrated confidence ordering rather than reliance on a single decision threshold.

From a methodological perspective, the modular two-stage design offers several advantages over end-to-end fine-tuning approaches. Separating representation learning from classification reduces sensitivity to proxy assay artifacts and facilitates reproducible benchmarking under distribution shift. Moreover, the framework is readily extensible: alternative genomic language models or additional feature modalities can be incorporated at the representation level without altering downstream task architecture.

Several limitations should be acknowledged. First, the model relies exclusively on sequence-derived features and does not incorporate epigenetic context, chromatin accessibility, or cell-type-specific information, all of which are known to influence Cas9 activity. Integrating such signals within a decoupled representation framework represents an important direction for future work. Second, the modest size of the TrueOT benchmark limits the statistical power of formal significance testing, highlighting the need for larger and more diverse experimentally validated datasets. Finally, the proxy training dataset excludes bulge-containing off-targets, constraining the scope of applicability to mismatch-based interactions.

Future work may explore controlled fine-tuning strategies for pretrained encoders, multi-task or multi-modal pretraining across genome-editing modalities, and the incorporation of chromatin-aware features while preserving strict external validation. As standardized benchmarks and evaluation protocols continue to mature, modular transfer-based frameworks such as the one presented here may serve as robust baselines for assessing generalization in CRISPR off-target prediction.

Overall, this study provides empirical evidence that decoupled representation learning can mitigate the limitations of proxy-trained models, offering a reproducible and extensible foundation for improving the reliability of computational CRISPR-Cas9 off-target prediction.

## 5 Conclusion

This study presents a modular two-stage framework for CRISPR-Cas9 off-target prediction that explicitly decouples sequence representation learning from downstream classification. By leveraging frozen pretrained genomic embeddings within a hybrid CNN-Transformer model trained solely on proxy assay data, the proposed approach improves generalization to experimentally validated off-target sites under strict external evaluation.

The results demonstrate that representation transfer provides a measurable benefit beyond architectural complexity, yielding improved discrimination, ranking quality, and confidence ordering on the TrueOT benchmark. Importantly, the framework emphasizes reproducibility and controlled evaluation under distribution shift, addressing a key limitation of many proxy-trained CRISPR scoring models.

Thus, this work highlights the value of transfer-aware modeling strategies for improving the reliability of computational off-target prediction. The proposed framework provides a reproducible and extensible baseline for future research and offers a principled foundation for integrating additional biological context as curated benchmarks and genomic resources continue to expand.

## Acknowledgements

The author thanks the open-source research community for publicly available datasets and tools that made this work possible. Computational resources provided by publicly accessible platforms are gratefully acknowledged.

## Declarations

### Ethics approval and consent to participate

Not applicable.

### Consent for publication

Not applicable.

### Data availability

This study uses two distinct datasets for training and evaluation:

**TrueOT Benchmark (Evaluation Dataset).** The TrueOT benchmark comprises 1903 experimentally validated CRISPR-Cas9 off-target sites curated by Kota et al. [10] from 11 low-throughput studies using techniques such as GUIDE-seq, CIRCLE-seq, and Digenome-seq. This dataset assesses model generalization from proxy training data to biologically validated off-target sites [10].

The processed file containing 1806 triplets was used for evaluation, it was derived from the original publication sources. Due to licensing constraints, this processed dataset is not redistributed. Users must download the original unprocessed data from: baolab-rice/CRISPR OT scoring TrueOT data

**Proxy Training Dataset.** Models are trained exclusively on a proxy dataset of 2,764 gRNA-target pairs curated from high-throughput GUIDE-seq assays by Park et al. [3]. Only mismatch-based off-targets were included (bulge-containing sequences excluded for architectural consistency). Labels were binarized using the original study’s cleavage efficiency threshold. This dataset is split into training/validation sets and never used for TrueOT evaluation.

**File Organization.** Place both processed datasets in the repository’s *data* directory following the provided preprocessing documentation. TrueOT data remains external (test-only) to ensure rigorous generalization assessment.

### Materials availability

Not applicable.

### Code availability

All code used for model implementation, training, evaluation, and figure generation is publicly available at: https://github.com/nyla-bhargava/Transformer-CNN-CRISPR

The repository includes:

- End-to-end training and evaluation scripts
- Deterministic random seed control
- Configuration files and hyperparameter specifications
- Instructions for reproducing all reported results

All experiments were conducted using fixed random seeds and deterministic data splits. Performance metrics reported in this paper were obtained using the exact evaluation scripts provided in the repository and were repeated with fixed preprocessing and evaluation protocols to ensure that observed performance trends are not artifacts of random initialization.

### Author contributions

NB conceived the study, designed the methodology, implemented the models, and drafted the manuscript. AG contributed to experimental design, result interpretation, and manuscript revision. All authors read and approved the final manuscript.

